# Characterization of walking locomotion in aged C57BL/10 mice: a comprehensive gait analysis

**DOI:** 10.1101/2023.04.04.535545

**Authors:** Joyce Santana Rizzi, Caique de Barros Almeida, Lorraine Silva Requena, Vitoria Duarte de Almeida, Thiago Vinícius Mininel, Gabriela Larissa Lima da Silva, Patrícia Fidelis de Oliveira, José de Anchieta de Castro e Horta Júnior, Cintia Yuri Matsumura, Renato Ferretti

**Author notes:** Corresponding Authors (RF). These authors contributed equally to this work (Joyce Santana Rizzi, Renato Ferretti). These authors also contributed equally to this work (Caique de Barros Almeida, Lorraine Silva Requena, Vitoria Duarte de Almeida, Thiago Vinícius Mininel, Gabriela Larissa Lima da Silva). These authors contributed equally to this work (Patrícia Fidelis de Oliveira, José de Anchieta de Castro e Horta Júnior, Cintia Yuri Matsumura).

## Abstract

Rodent gait analysis is crucial for modeling human aging, but the lack of comprehensive research on gait in elderly mice limits our ability to translate findings from animal models to human populations. Age-related changes in C57BL/10 strain remain unknown. The state of art protocol for gait analysis uses the CatWalk ™ XT system that allows an understanding of the locomotion pattern by a variety of parameters. We aim to provide relevant information for experimental designs, presenting benchmark data on the performance of locomotion using healthy wild-type mice for future preclinical investigations of neurological and neuromuscular gait patterns. In this study, characterization of walking locomotion was demonstrated from complete gait analysis in aged C57BL/10ScCr/PasUnib mice using open-field, CatWalk, and treadmill tests. Mice were divided into the adult group (6 months; n = 9) and the aged group (20 months; n = 9). Aged mice demonstrated decreased mobility, distance traveled, and general speed in the open-field test. The spatiotemporal and kinetic parameters were altered in aged mice, with lower speed, higher stand time and stride length, and increased base of support and duty cycle in comparison with adult mice. Interlimb coordination has changed in elderly mice. To test whether speed alters the temporal parameters, we used a treadmill test and we demonstrated higher stand time in 20-month-old mice. We demonstrated that changes in gait parameters and mobility represent direct age-related singularities in the wild-type C57BL/10 mice. Overall, aged mice took more time in contact with the ground independently of the speed. These baseline gait results shed light on measures that allow the potential investigation of therapeutics and interventions in gerontology or neuromuscular diseases.

## Introduction

Aging affects many physiological systems of the body due to impairment in the nervous and musculoskeletal systems, which can increase fear or the risk of falls in old age [1,2]. Walking performance is largely influenced by the decrease in muscle strength and coordination, and older populations tend to exhibit a more cautious gait characterized by alteration of spatial-temporal parameters (slower walking speed, shorter stride length, and increased step width) [2,3]. A decrease in walking performance, especially walking speed, is a hallmark of health, as it predicts mobility independence, cognitive function, survival rate, and mortality [4]. Various assessments of gait parameters provide more sensitive and quantitative measures of motor control in experimental models [5-10]. Gait analyses are often used to test motor and coordination dysfunction based on diagnostic criteria for many neurological and neuromuscular disorders [11,12]. Researchers used spatiotemporal parameters as control values to determine if a particular population deviates from the norm, for example, when investigating the influence of neurological, cardiovascular, or musculoskeletal impairments on gait [1,3,5,7,8]. However, gait qualitative and quantitative measurements are inconsistent and the age-related changes in gait parameters remain unknown in C57BL/10 mice.

The CatWalk system is widely used as an automated and computerized gait analysis system that utilizes a walkway glass plate to capture digital images of animal paws as they move through the platform [6]. This system is superior to other technologies in providing high-quality images of paw abnormalities and providing observed-independent quantitative assessments of spatiotemporal, kinetic, and coordination parameters [6]. Despite the differences between quadrupedal and bipedal gait patterns, basic data collection methods remain analogous [4,7]. Many genetic, pharmacological, and nutritional interventions have been shown to increase the lifespan of laboratory mice [13-17]. Preliminary evidence suggests that mouse gait is altered in pathophysiological conditions associated with accelerated nervous system aging [5-9,13]. Despite the widespread use of the C57BL/6 mice to identify translational preclinical measurements of physiological dysfunction [4-12], the C57Bl/10 mice model remains concerned regarding their gait validity.

In the present study, we conducted a comprehensive gait analysis in aged C57BL/10 mice to establish benchmark data for preclinical investigation. The walking locomotion characterization of the C57BL/10 strain was demonstrated by performing a complete gait analysis in aged mice using open-field and CatWalk system. We assessed whether constant speed in treadmill test alters temporal gait parameters. Our results demonstrated that aging reduces walking speed and stride length while increases stand time, the base of support, and limb support. By comparing gait function in freely moving 6- and 20-month-old mice using the computer-aided system for the quantification assessment of footfall and motor performance this present study further expands our understanding of the impact of aging on gait pattern. Also this provides useful insights for future preclinical approaches to study circuits in abnormal gaits experimental models, such as neurological and neuromuscular diseases.

## Materials and methods

### Care of animals and experimental design

Eighteen male wild-type mice (C57BL/10ScCr/PasUnib) were obtained from the breeding colony at the State University of Campinas (CEMIB-UNICAMP) and maintained by our institutional animal care facility. All animals were divided into two groups: adult mice (6 months; n = 9) and aged mice (20 months; n = 9). Three animals were kept in cages with enrichment materials (cardboard tunnels and sheltering device) at constant temperature (21 ± 2°C), cycles of 12h light/ 12h darkness (lights 7 a.m. - 7 p.m.), and with free access to food and water. All tests were conducted during the light phase (between 10 a.m. and 4 p.m.). The experimental procedures were performed in accordance with the Brazilian Guide for Care and Use of Laboratory Animals and approved by the Animal Care and Use Committee (process number 1401/2021). The test order started with the open-field test provided by a 2 hours room habituation; the CatWalk tests were performed after three consecutive days of adaptation; and a week after, all mice were adapted for 3 minutes and submitted to the treadmill test.

### The open-field experiment

Adult and aged mice (n=9 per group) were tested in the open-field experiment. A circular open field arena (circumference of 43 cm, height of 25 cm) was painted white and the floor was divided into 2 black circles (6 cm ratio and 23 cm ratio), subdivided by black lines ruled on the ground that formed 18 trapezoid spaces. All mice were acclimated in the room for 2 hours before the test. At the start of each test period, the mouse was placed gently in the middle of the open field for one 5-min period (between 1:00 pm and 3:00 pm). A high-speed video camera (100 frames/s) was positioned 50 cm above the open field, and the tracking video was analyzed using the Image J software. At the end of each test period, the mouse was removed, placed back in the original cage, and the open field was sanitized with 45% ethanol. We measured the distance traveled; the percentage of the time of immobilization (defined as no movement for more than 2 seconds), and the average speed collected and analyzed after all mice completed the test.

### The CatWalk™ gait analysis system

After the open-field test, all mice were habituated to the experimental procedures for three consecutive days. A single trained operator performed gait experiments on the fourth day to avoid bias. To motivate the animals to cross the glass plate we attached the home cage to the end of the CatWalk walkway. The apparatus was cleaned with 75% ethanol solution and dried with a paper towel in between each mouse. No additional food or other rewards were used in our experiments. The CatWalk™ XT analysis system (Noldus, Netherlands) consisted of a glass platform with a high-speed infrared camera (100 frames per second), and CatWalk™ software for tracking and analyzing free locomotion. Mice were placed in the catwalk platform for a short time (3-5 minutes) to get them used to the environment and then free locomotion was recorded for a maximum of 10 minutes. A valid walking test was performed when the animals walk without refusal, pause, or jumps. Three complete stride cycles were considered a validated walk and the average was used to compare adult and aged mice. The CatWalk software collected all parameters automatically and the trained operator conducted the experiments. The data on the gait parameters were divided into six categories: general characteristics, spatial, temporal, kinetic, interlimb coordination, and intensity-related parameters (Table 1). Each parameter provided by the CatWalk™ XT system allows the understanding of the gait pattern through the evaluation of the categories (S1-Table1). The general characteristic parameters describe the cadence (steps per second), number of steps, duty cycle (ratio of stand time to step cycle), and the base of support (width on the y-axis between either forepaws or hindpaws). The spatial category describes the area, distance, length, and width of the paws or stride, represented in centimeters or area (cm^2^). The temporal category is represented in time, its parameters indicate the time necessary to execute a certain stage of the stride, usually, the data represented are described in seconds or percentages. The kinetic category describes parameters that indicate movement, and describe the relationship between distance and time during stride stages. Interlimb coordination indicates how the paws were distributed along the stride cycles, usually indicated in percentage and arbitrary units. Finally, the intensity-related parameters indicate the relationship between the force applied and distributed on the paws during the steps of the stride cycles, which can also be related to time. These are usually represented in percentages or arbitrary units. In our study, we kept a constant gain of 37.9 and an intensity of 0.17 between each mouse.

### The treadmill gait analysis

One week after the CatWalk protocol, all mice were exposed to the treadmill protocol. The treadmill (AVS, Sao Carlos, Brazil) was placed on a level surface and a high-speed camera (65 frames/s) was positioned 10 cm distance from the belt to capture the gait cycle from a lateral view. A range of running speeds was tested in a pilot experiment to define an optimal speed that mice were able to run independently of their ages, thus excluding differences in self-selected speeds as the critical confounding factor in the interpretation of gait analysis. The mice were gently placed on the stopped treadmill belt for 3 minutes to adapt and then induced to walk on the treadmill at constant 10 cm/s for a maximum of 5 minutes. Three complete gait cycles were recorded. The walk without refusal, pause, or jumps was considered valid. Image J motion analysis plugin software was used to process the video to determine the spatial gait parameter, such as stride time, stance time, and swing time. The stance time was subdivided into brake time and propulsion time.

### Statistical analysis

All statistical analyses were performed using GraphPad Software (La Jolla, California, USA, www.graphpad.com). Three runs were tested by two-way ANOVA followed by Bonferroni correlation and the average was used for the comparison of aging. For the comparison of adult and aged mice, we used the unpaired Student *t*-test with Welch’s correction, assuming different standard deviations. The confidence level was 95%, and the definition of statistical significance was p ≤ 0.05. Ordinary two-way ANOVA followed by Bonferroni’s post hoc test for multiple comparisons to determine differences between ages and paws. Significance was considered as p ≤ 0.05.

## Results

### Open-field test

Spontaneous locomotion was measured in all mice using the open-field test to detect age-related changes in general motor activity. Gait parameters were evaluated in free movement using the CatWalk system, and to test the influence of speed on temporal gait parameters we assessed a constant speed test using the treadmill. The distance traveled in the open-field test by aged mice was significantly lower than by adult mice (Fig 1C). Aged mice had a higher inability to free movement and immobility time was greater than adult mice (Fig 1D). During five minutes of free locomotion, aged mice exhibited a reduced average speed (Fig 1E) compared to adult mice.

**Figure 1.**
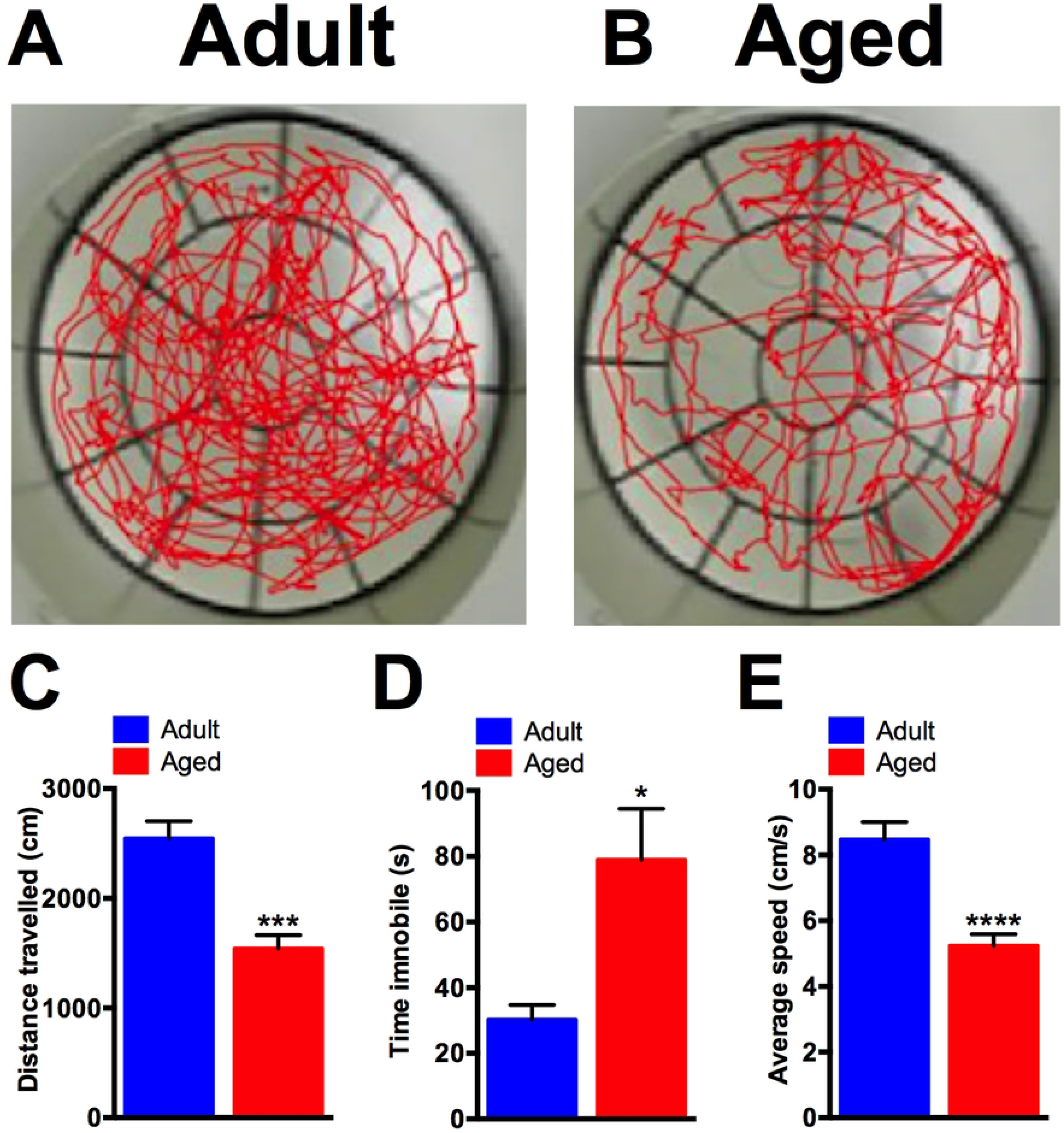
Spontaneous locomotion in open-field test in aged mice. A-B. Tracking movement in the open-field test between 6-month-old mice (Adult) and 20-month-old mice (Aged). C. Response of the age to the traveled distance in comparison to adult mice. D. The effect of aging immobility. E. Decrease in average speed between adult and aged mice. n = nine mice in each group; * p< 0.05; *** p< 0.001; **** p< 0.0001.

### The CatWalk test

The general characteristics and interlimb coordination parameters used in CatWalk gait analysis are commonly utilized to identify neurological and neuromuscular abnormalities. Our study showed that certain characteristics and interlimb coordination slightly change in 20-month-old mice (Table 1). The percentage of stand over the sum of stance and swing duration increased substantially when mice get older (Duty cycle). The cadence, number of steps, and regularity index (a coordination parameter) did not vary with aging. However, aged mice exhibited an increase in the base of support, and a more predominant alternate footfall pattern AB (i.e. a normal sequence LF-RH-RF-LH). During free locomotion, aged mice spent more time standing on diagonal, lateral, and three support paws compared to adult mice, which primarily used a single paw support.

To determine the interaction between age and paws in the gait parameters we conducted a two-way ANOVA test, and we found that age was significantly different rather than the involvement of each paw (S1-Table 2). Since aging has a major impact on these parameters, we represented our data using the mean of paws and we divided the gait parameters into categories (as outlined in Table 1). Although the total walk duration and step time were similar between ages, 20-month-old mice took approximately 9% more time during the stance phase, which resulted in an increased stand time while freely walking. Aged mice decrease speed and stride length compared to adult mice. Aged mice also kept a single paw in contact with the ground for a longer period than adult mice; particularly their hind paws (S1-Table 2). The stride length of limbs decreased significantly, whereas the footprint length, width, and area increased substantially. The step length measured by the print position was comparable between ages. At the kinetic parameters, 20-month-old mice had a significantly decreased in walk speed, a decrease in swing speed, and an increased speed at which the paw lost contact with the ground (i.e. the stand index). A two-way ANOVA particularly revealed an interaction between ages in front and hind paws on swing speed (F_1.32_ = 4.32; p=0.04), and between ages in stand index (F_1.32_ = 6.516; p=0.01).

The camera gain and light intensity on the CatWalk can impact the intensity-related parameters. In our study, we utilized a constant gain of 37.9 and an intensity of 0.17 between each mouse (as described in the Material and Methods section). Our finding indicated that the maximum contact mean intensity, and the minimum and mean intensity of a complete paw were reduced in aged mice. No significant differences were observed in the other intensity-related parameters during the aging process.

**Table 1. Gait abnormalities in aged mice.** The mean and standard error measures of CatWalk parameters from adult and aged C57BL/10 mice. Each gait parameter was categorized. For the comparison of adult and aged mice, we used the unpaired Student *t*-test with Welch’s correction, assuming different standard deviations. The confidence level was 95%, and the definition of statistical significance was p ≤ 0.05. ns = no statistical significance; * p< 0.05; ** p< 0.01.

### The treadmill test

To address the issue of speed dependence in the gait parameters, we performed a gait analysis at a constant speed using a treadmill. Our results showed that the stride time was higher in aged compared to adult mice (Fig 2A). At an equivalent speed, the stance time (Fig 2B) was higher in aged compared to adult mice; while the swing time was similar between ages (Fig 2B). We subdivided the stance phase into brake (initial contact of the paw to full paw contact) and propulsion (full paw contact to final contact) phases. 20-month-old mice seem to spend more time in propulsion (Fig 2D) than in the brake phase (Fig 2C) compared with adult mice. Besides speed, aged mice took more time in the stance phase than the swing phase, resulting in an increased stance time and reduced swing time.

**Figure 2.**
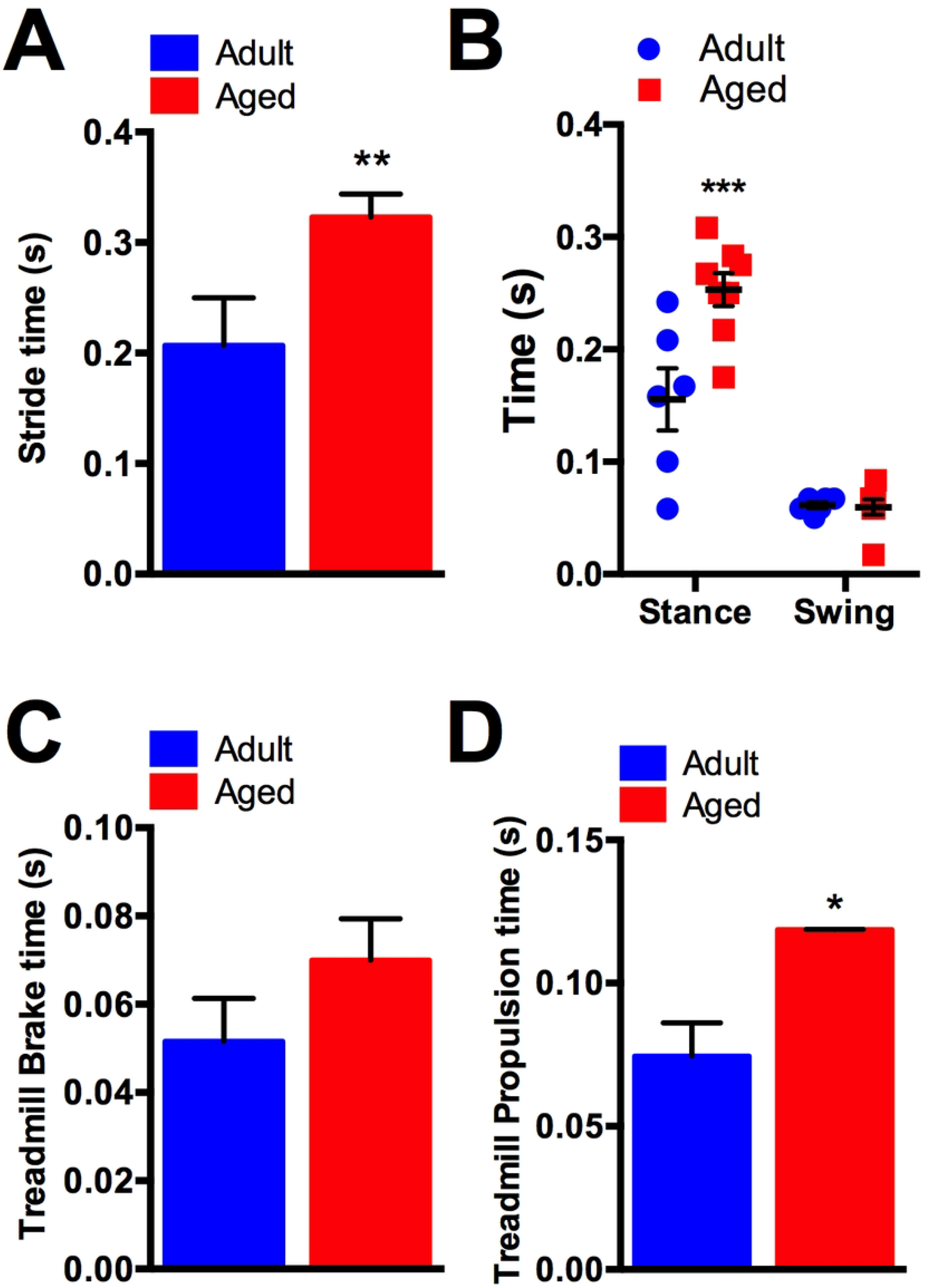
Age-related changes in gait analysis at a constant speed during a treadmill test. A. Increased stride time during treadmill test between 6-month-old mice (Adult) and 20-month-old mice (Aged). B. The effect of aging on the mouse’s time of stance and swing phase, respectively. C. Subdivision of stance phase into brake time, and D. propulsion time. n = nine mice in each group; ** p< 0.01; *** p< 0.001.

## Discussion

Gait analysis is a tool for evaluating motor function in aging and neuromuscular diseases. We demonstrated aging-induced abnormalities in locomotion and gait parameters in a commonly used wild-type mouse model. In the open field test, gait parameters and general locomotion were studied in 20-month-old C57BL/10 mice that exhibited decreased average speed, mobilization, and distance traveled. Using the CatWalk system, aged mice demonstrated variability in the spatiotemporal, kinetic, interlimb coordination, and intensity-related parameters compared with 6-month-old mice. Finally, at a constant speed, we observed an increase in the stance time in aged mice. Overall, our results indicated that the aged mice took more time in contact with the ground independently of the speed.

The open field test is a friendly method used to evaluate behavior and demonstrates locomotion impairments. We exposed all animals once in their lifetime to the open field test to study spontaneous and general locomotion. Aged mice reported significant locomotion impairments with shorter total distance traveled, lower average speed, and longer period of immobilization than 6-month-old adult mice. Even in healthy conditions, the decline in walking speed and mobility is, to some degree, a consequence of normal aging [18, 19]. Walking is an indicator of overall health, and walking speed is correlated with life span expectancy [18]. Aged C57BL/10 mice have shown drastic locomotion impairments in the open-field test compared to C57BL/6 mice [19]. The results of this study generally support previous findings that locomotion activity decreases with increasing age [18, 19]. However, the cause of the decreased walking performance and cognitive function in C57BL/10 is still not fully understood.

Aging resulted in a shorter stride length, larger paw area, and lower walk speed [18, 19]. The increased paw contact area may suggest incomplete or ineffective foot lift of the ground during locomotion, causing an increase in total foot contact with the ground instead of just the toe contact [20]. Our open-field experiment further support the impact of aging on behavior and mobility, revealing a significant reduction in the ability of aged mice to move, covering a shorter distance and exhibiting a lower speed compared to adult mice. However, gait parameters are not fully known in C57BL/10 mice. We did not analyzed gait parameters in open-field test but all mice were adapted to walk spontaneously in the CatWalk system.

To investigate age-related changes in gait parameters, we used the CatWalk analysis system and a treadmill. The CatWalk system is mostly used equipment to assess gait patterns in rodents, and data collection represents static and dynamic parameters of free walking variability that were collected simultaneously [7, 20]. Gait abnormalities can be assessed using spatiotemporal and kinetic measurements, such as stride length, walking duration, cadence, stand and swing time, step cycle, duty cycle, and swing speed. The base of support, initial dual stance, diagonal support, and three or four support can identify posture alterations.

The age-related shifts in C57BL/10 gait parameters are distinct from those seen in different strains, ages, and sexes [20]. In our study, male 20-month-old C57BL/10 mice showed decreased walking speed, swing speed, and stride length, while the stand time, duty cycle, and base of support were increased compared to 6-month-old mice. However, the elderly 24-month-old C57BL/6 mice did not show a statistical difference in walking speed, stride time and length, swing speed, duty cycle, or the base of support compared to 3-month-old mice [19]. Females were affected earlier than males C57BL/6 with greater stride length, swing, and support time, and decreased cadence as early as 23 months [22]. However, we did not compared sexes interaction between gait parameters within the same age. Further analyses are needed to investigate whether female C57BL/10 mice shift gait marks earlier than male.

Different mice strains have been used in aging research to identify key mechanisms to test life-span compounds. In the rotarod and the horizontal bar tests, the C57BL/10 fell off more frequently than C57BL/6 mice, while the maximum speed was similar between strains, which indicate that the C57BL/10 mice have coordination impairments rather than low strength [22]. Although C57BL/10 and C57BL/6 mice share the same origin, there are phenotypic and genotypic differences between them. The C57BL/10 mice performed worse than C57BL/6 mice in behavioral tests, cognitive tests, and motor tests [22]. Locomotion abnormalities were linked to sensorimotor integration, coordination, and nervous system function, which can lead to a more cautious gait and a decrease in walking speed. Nevertheless, the trajectory of change-driving aging in C57BL/10 differs from those in C57BL/6 mice.

The pattern of age-related shifts in gait parameters observed in this study diverges from the pattern seen in later stages of wild-type mice [21] and in experimental models of Parkinsonism or ataxia [23]. 20-month old C57BL/10 mice presented decreased stride length and speed, while cadence was similar to 6 month-old mice. Age-related gait changes may be affected through different neural circuits, from the caudal brainstem to the upstream central nervous system. Aging processes change temporal gait parameters and seem to be associated with walking speed [21]. Despite the assertion that gait parameters stand if there is no alteration in speed from stride to stride, we submitted all mice to a second round of walking with a constant velocity on a treadmill. We found a decrease in the temporal parameters in aging mice. In contrast to humans, the mice are not able to maintain a steady speed or modulate velocity between trials, leading to a huge challenge in experimental studies. However, we adapted all mice to the treadmill and used a 10 cm/s speed to allow the animal to walk freely. The walking speed declined earlier during the aging process than stride length and swing time [21]. Furthermore, in humans, the decline in speed is caused by a decrease in stride length rather than by a change in cadence [24]. Furthermore, aged mice had similar steps per second (cadence) compared to adult mice. Nonetheless, using a treadmill for gait analysis did not eliminate the speed-related issue, as it precludes the measurement of the preferred speed, which may be a characteristic of that particular group or strain.

The single fibers in aged C57BL/10 mice differentially presented both type I and II myosin heavy chains (MyHC), increased resistance in isometric fatigue, lower tension, and shorter speed compared with young ages [25]. Aging promote progressive impairment in muscle mass and strength, which seem to be mediated by mitochondria dysfunction [26]. Nonetheless, changes in muscle contractile properties and mitochondria function during the aging process in C57BL/10 mice are still unknown. Hind-limb gait parameters reveal striking similarities among mammals [4,20]. Despite the differences in size and musculoskeletal anatomy among species, the similarities in hind paw patterns are noteworthy [4]. Although our results did not present paw interaction in the majority of gait parameters, the temporal parameters in hind paws have decreased compared to front paws (S1 Table 2). Thus, some gait parameters seem to be affected by dominant paw or wild-type strain.

Although the C57BL/10 strain is widely used for a variety number of diseases models [27-33], this mouse strain is utilized as a reference for comparisons with the *mdx* mice, the widely used model of Duchenne Muscular Dystrophy (DMD) [34]. DMD is a progressive neuromuscular disorder caused by mutations in the dystrophin gene, characterized by muscle degeneration and weakness, and leading to premature death from respiratory failure or cardiac dysfunction [35]. Our findings reflect alterations in gait function in 20-month-aged wild-type mice that provide important reference values for the design of future studies attempting to characterize the effects of dietary, pharmacological, or genetic interventions on health and neuromuscular conditions, such as Duchenne Muscular Dystrophy.

Our study provided baseline parameters of C57BL/10 mice and highlighted the need for a comprehensive analysis of aging in each strain to increase the reliability of mice as models of human aging. We conducted a comprehensive gait analysis in aged C57BL/10ScCr/PasUnib mice to establish benchmark data for future preclinical investigations. Furthermore, the C57BL/10 gait signature demonstrates that the wild-type mouse database is an important preclinical tool for investigating health and disease conditions. Future studies are necessary to fully comprehend the impact of aging on locomotion in different strains, ages, and sexes, and the development of strategies to improve mobility and quality of life in aging populations. Furthermore, locomotion decreases is a sensitive indicator, which in older humans was reported to predict cognitive decline and decreased survival, and is associated with the risk of falls.

## Conclusion

We demonstrated that changes in gait parameters and mobility represent direct age-related singularities in the wild-type strain. These results established the baseline of gait data for widely used wild-type C57BL/10 mice, which allows for its comparison against treatment groups and the potentially meaningful investigation of translatable therapeutics and interventions. Also, this study shed light on the necessity to carefully select the parameters of outcome measures in automatic or interventional gait studies.

## Acknowledgments

Bruno dos Santos Ricardi, Diego Generoso, and Vitória Luiza Alves de Souza Rios for their technical assistance in the CatWalk system and their valuable comments on the manuscript.

## Support information

**S1 Table 1**. Each parameter provided by the CatWalkTM XT system allows the understanding of the gait pattern through the evaluation of the categories.

**S1 Table 2**. Age was significantly different rather than the involvement of each paw.

